# The olivary input to the cerebellum dissociates sensory events from movement plans

**DOI:** 10.1101/2022.07.11.499614

**Authors:** Jay S. Pi, Mohammad Amin Fakharian, Paul Hage, Ehsan Sedaghat-Nejad, Salomon Z. Muller, Reza Shadmehr

## Abstract

Neurons in the inferior olive are thought to anatomically organize the Purkinje cells (P-cells) of the cerebellum into computational modules. To better understand what is computed by these modules, we designed a saccade task in marmosets that dissociated sensory and motor events and then recorded the complex and simple spikes of hundreds of P-cells. We found that when a visual target was presented at a random location, the olive reported the direction of that sensory event to one group of P-cells, but not to a second group. However, just before movement onset it reported the direction of the planned movement to both groups, even if that movement was not toward the target. At the end of the movement if there was an error but the subject chose to withhold the corrective movement, only the first group received information about the sensory prediction error. We organized the P-cells based on the information content of their olivary input and found that in the group that received sensory information, the simple spikes were suppressed during fixation, then produced a burst before saccade onset in a direction consistent with assisting the movement. In the second group the simple spikes were not suppressed during fixation but burst near saccade deceleration in a direction consistent with stopping the movement. Thus, the olive differentiated the P-cells based on whether they would receive sensory or motor information, and this defined their contributions to control of movements as well as holding still.

**Significance statement:** We found that in the oculomotor region of the vermis, the olive informed a subset of P-cells about sensory events that were prediction errors, but all P-cells about the forthcoming movement. Using the information content of the olivary input we labeled the P-cells and produced a population code, revealing two groups with simple spikes that were antagonistic to each other, one contributing to movement initiation, the other signaling movement end.

## Introduction

Theories of cerebellar function suggest that we learn how to move because the inferior olive provides a teaching signal to the Purkinje cells (P-cells) of the cerebellum (1–3). In this framework, the resulting complex spikes (CS) encode an error (3–5) that trains the simple spikes (SS) (3, 6), the output of P-cells. However, the occurrence of an error is not the only event that modulates the CS rates (7): while the rates are modulated following a variety of sensory events (8–13), they are also modulated before a variety of movements that are voluntary (14–18) and thus are not errors. For example, the CS rates are modulated by sudden sensory events such as the touch of a limb (19), an air puff to the face (10, 11, 20), or presentation of a visual stimulus (9, 21), signaling that an unpredicted sensory event has occurred (10, 13), particularly one that is salient (12). But the CS rates are also modulated before voluntary movements such as walking (16), licking (14), reaching (8, 15, 17, 18, 22), and moving the wrist (23), none of which are unexpected or erroneous. This diversity of events that affect the CS rates begs the question: what information is encoded in the climbing fiber input to the cerebellum? Inspired by the behavioral experiments of Wallman and Fuchs (24), and Tseng et al. (25), we designed a 2x2 saccade task in which a visual event was sometimes but not always followed by a movement, and a movement that was sometimes but not always preceded by a visual event. At the conclusion of some movements, the subject experienced an error. We used reward to assign a value to this error, affording the subject the opportunity to respond to that error by making a corrective movement, or simply observing the error but withholding the movement. This allowed us to quantify the CS response to visual events, motor events, and sensory prediction errors that did or did not produce corrective actions.

We recorded spiking activities of hundreds of P-cells in the oculomotor vermis of marmosets and found that following the presentation of a visual target at a random location, the olive reported the direction of that target at low latency (60 ms), and then just before the ensuing movement, it reported the direction of the planned saccade, even when that saccade was not toward the target. However, the olive differentiated the P-cells into two populations: in one group (Type 1), the P-cells received information regarding the direction of the target, and then independently, the direction of the planned movement. In another group (Type 2) the P-cells received information regarding the planned movement, but not the visual event.

At the conclusion of some saccades the target was not on the fovea, indicating a sensory prediction error. On trials in which the subject chose to make a corrective movement the CS response in both groups of P-cells encoded the error direction. However, if the subject chose to withhold the movement, the error information was still transmitted from the olive to the P-cells, but only to the Type 1 group. That is, the olive reported sensory prediction errors to only the Type 1 P-cells. Notably, the olive only weakly reported sensory prediction errors that did not inspire corrective movements.

It is possible that two groups of P-cells that receive different kinds of information from the olive during a given behavior contribute differently to control of that behavior (26). Indeed, the P-cells whose olivary input encoded the visual target had an SS response that acted as an agonist, promoting the movement toward the target. In contrast, the P-cells whose olivary input encoded the forthcoming movement but not the visual target had an SS response that acted as an antagonist, resisting the movement as it neared the target.

## Results

Marmosets fixated at a center location and were presented with a primary target at a random direction (Fig. 1A). As the primary saccade commenced, we moved the target to a random secondary location, thus inducing a sensory prediction error that was usually corrected by a movement (Fig. 1B). To assign a value to this error, in 60% of the sessions reward required both the primary and the secondary saccades (Fig. 1A, left subplot), whereas in the remaining sessions reward required only the primary saccade (Fig. 1A, right subplot). If reward did not require the secondary saccade, the secondary target was presented briefly (∼120 ms, adjusted during the recording session) and then returned to the primary location. In this case, the primary saccade ended with a sensory prediction error, but one that did not produce a corrective movement. Following fixation of the final target, the center target was presented, and the subject made a center saccade. Thus, the 2x2 design (Fig. 1C) allowed us to differentiate between the responses to visual events, motor events, and sensory prediction errors that followed a movement but did or did not produce a corrective action.

**Figure 1.**
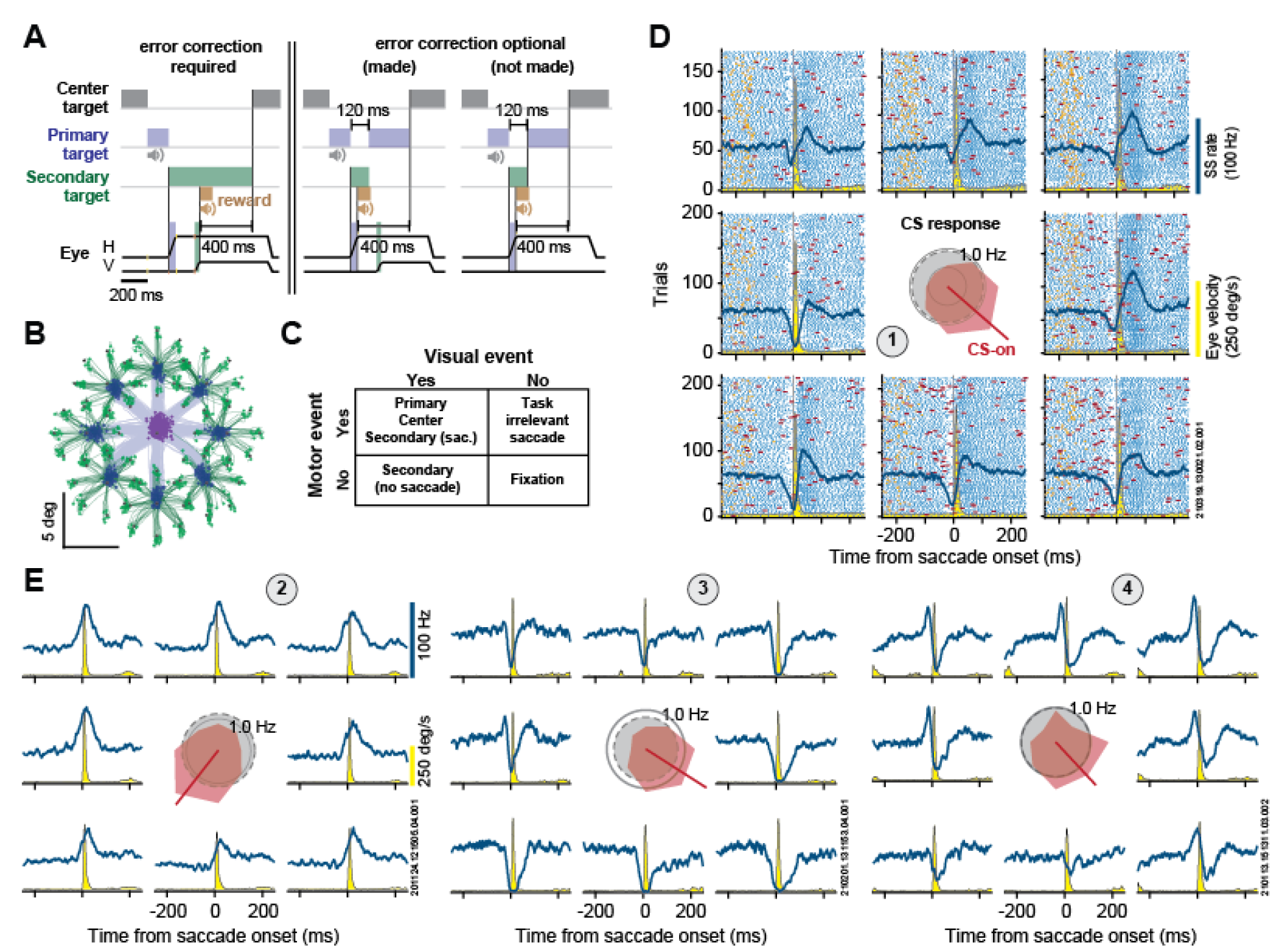
A saccade task designed to dissociate sensory and motor events. **A**. Task design. In some sessions the primary saccade concluded with an error and reward required correction via a secondary saccade. In other sessions this correction was optional as the reward was provided with the primary saccade. **B**. Saccade trajectories for primary (blue) and secondary (green) saccades. **C**. 2x2 design. Some but not all visual events were followed by a movement, and some but not all movements were preceded by a visual event. **D**. Spiking of an example P-cell. CSs (red) and SSs (blue) of a P-cell are aligned to the onset of the primary saccade. The average CS rates before saccade onset as a function of direction is shown at center. Gray circle in the center plot indicates baseline CS rates, computing as the average CS rate in the entire recording. The yellow trace is saccade velocity. **E**. Example of P-cells with similar CS tuning (cells 1, 3, and 4) but rather different SS patterns.

We combined MRI and CT guided targeting procedures (27) to place tetrodes, heptodes, and silicon probes in lobules VI and VII of the vermis and recorded from 381 P-cells over a 2.5 year period in 2 marmosets. In every case, the neuron was identified as a P-cell because of the presence of CSs. In 254 P-cells we isolated both the SSs and the CSs and confirmed that the CS suppressed the SS. The data set for each P-cell included an average of 1530±586 (mean ± median absolute deviation) task-relevant saccades to visual targets and an additional 1618±908 task irrelevant saccades.

In general, the SS response, as exemplified in the four P-cells shown in Figs. 1D & 1E, varied widely between the cells (5, 28, 29). For example, during the primary saccade, cell 1 exhibited a pause-burst pattern, cell 2 exhibited a burst pattern, cell 3 exhibited a pause pattern, and cell 4 exhibited a burst-pause pattern. However, despite these diverse SS modulations, cells 1, 3, and 4 all had their largest CS response before saccades in direction -45° (termed CS-on, (9)). We used the CS-on direction of a P-cell to define a coordinate system for that cell (5, 6, 30). As a result, for cells 1, 3, and 4, the saccade toward 0° was labeled as CS+45°, whereas the same saccade for cell 2 was labeled as CS+135°.

### The olivary input reported the direction of the visual target as well as the direction of the forthcoming saccade

We measured the CS response of each P-cell following the presentation of the primary target and labeled the direction that produced the largest rate increase as CS-on (Supplementary Fig. S1A). The CS- on direction remained consistent regardless of whether we measured it in response to the primary target, the secondary target, or the center target (Supplementary Fig. S1C). Notably, the CS-on direction varied approximately with the cell’s location in the vermis (Supplementary Fig. S1B) (9, 30). P-cells on the right side of the vermis tended to have their largest CS response to visual events to the left of fovea, whereas the P-cells on the left side produced a CS in response to events on the right.

When the target was in direction CS-on of a P-cell, that cell responded with an increased CS rate, but with a timing that differed among the cells (Fig. 2A, middle row). For each cell we measured the timing of the peak CS response, labeled it with a random variable 𝑌, and found that in the population, the distribution of 𝑌 appeared bimodal (Fig. 2A, top row). As a result, the CS firing rates also exhibited a double-peaked response, which was particularly prominent when we separated the trials based on reaction times (RT) (Fig. 2B, right column, the trials were divided based on the median of the RT distribution, 166 ms, mean: 195 ms). Moreover, when we aligned the data to saccade onset, the rates peaked at -10±1 ms (mean±SEM) before saccade onset (when the saccade was in direction CS-on). Thus, the CS rates appeared to be modulated both in response to the target, and in anticipation of a forthcoming movement.

**Figure 2.**
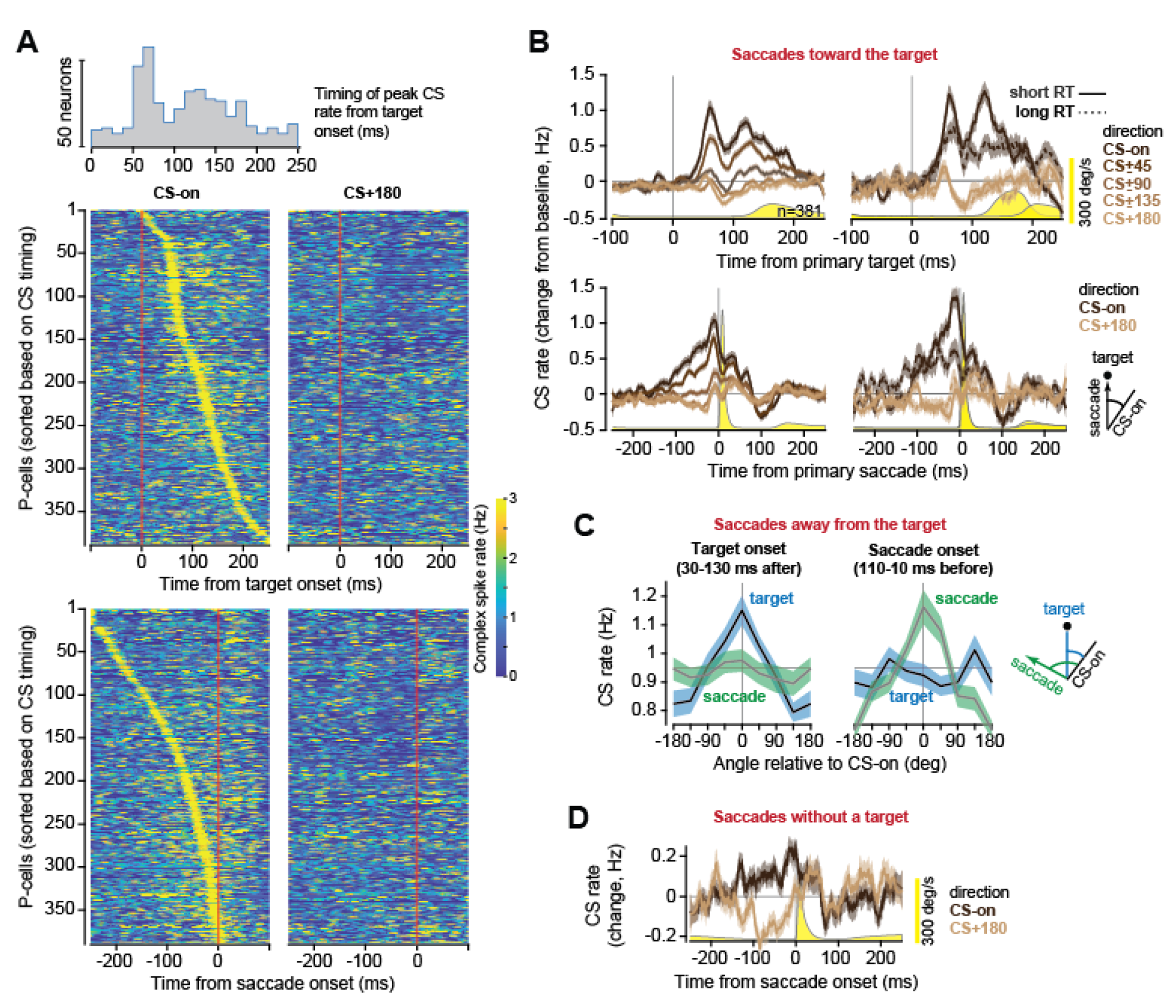
The olivary input independently reported the direction of the visual target and the direction of the forthcoming movement. **A**. CS response of all P-cells. The distribution is the timing of the peak CS rate from target onset. Cells were sorted based on timing of peak response following target onset, and before saccade onset. **B**. CS response of the population following the onset of the primary target, and before the onset of the primary saccade. The directions of the visual and motor events are with respect to CS-on. The plot in the right column illustrates the CS response in trials that exhibited short or long reaction times (RT). Yellow trace is saccade velocity. **C**. In some trials the target was presented but the subject chose to saccade elsewhere. The CS response following target onset encoded target direction, but the CS response before movement onset encoded saccade direction. **D**. Some saccades were not preceded by the onset of a visual stimulus and were reward irrelevant. Nevertheless, the CS rates were weakly modulated before these saccades as a function of the direction of the forthcoming movement. Error bars are SEM.

To dissociate target direction from saccade direction, we focused on trials in which following target presentation the subject chose to saccade elsewhere (defined as saccades not directed to within 45° of the target). For this analysis, we combined the saccades that were made in response to either the primary or the secondary target. The time between target onset and saccade onset for these saccades was 350±360 ms (mean±SEM), long enough to dissociate the CS response to the visual and motor events. We found that following target onset, the CS rates (30-130 ms after onset) encoded target direction, not the direction of the forthcoming saccade (Fig. 2C, average firing rate in direction CS-on vs. CS-off; Wilcoxon signed rank test; target: z=6.39 p=2x10^-10^; saccade: z=0.12 p=0.9). However, in the period before saccade onset (110-10 ms before), the CS rates encoded direction of the forthcoming saccade, not the direction of the target (Fig. 2C, average firing rate in direction CS-on vs. CS-off; Wilcoxon signed rank; target: z=0.4 p=0.67; saccade: z=7.97 p=2x10^-15^). Thus, we uncovered a double dissociation: the CS rates following target onset signaled target direction but not saccade direction. However, the CS rates before saccade onset signaled saccade direction but not target direction.

We next asked whether the olivary input reported the direction of the saccade even if that movement was not in response to a visual target. In task-irrelevant saccades, the movement was performed outside of the context of the task: the subject was not fixating at the center target, and there were no sudden or unexpected sensory events that we know of that inspired these movements.

Regardless, the CS rates were modulated before the saccades and carried information regarding the direction of the movement (Fig. 2D, CS-on vs. CS-off, 50 ms period before saccade onset, Wilcoxon signed rank test, z=6.59 p<10^-6^). However, the CS rates before these movements were much less modulated as a function of direction than for task relevant saccades (vs. primary saccade, 50 ms period before saccade onset, Wilcoxon signed rank test, z= 8.04 p<10^-6^).

In summary, the inferior olive provided the oculomotor vermis with two kinds of information: at short latency, it reported the direction of the random visual target, and then before the movement, it reported the direction of the forthcoming saccade, even when that saccade was not toward the target.

### The olivary input divided the P-cells into two groups

The timing of the CS response following target onset (Fig. 2A, 1^st^ row) suggested the presence of two groups of neurons: one group that responded with a CS at low latency, potentially encoding the direction of the visual stimulus, and another that responded with long latency, potentially encoding the direction of the planned movement. As a result, when the target was in direction CS-on, across the population the CS firing rate was double-peaked (Fig. 2B, first row), with the first peak at 65±2 ms (mean±SEM). We represented the CS timing distribution (Supplementary Fig. S2A) as a mixture of gamma functions and performed statistical testing to ask whether the distribution was composed of a mixture of 1, 2, or 3 groups of neurons.

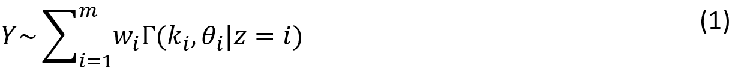

Both AIC and BIC suggested that the population was best represented as a mixture of two groups of neurons (Supplementary Fig. S2B, 𝑚 = 2, 𝑤_1_ = 0.68, 𝑘_1_ = 2.38, 𝜃_1_ = 17.9, 𝑤_2_ = 0.32, 𝑘_2_ = 39.1, 𝜃_2_ = 9.3). Because in the population the two peaks in the CS response had a minimum that occurred at ∼100 ms, we divided the population based on the criterion of whether the CS response had its maximum peak during the period of 0-100 ms, or after this period. We labeled the P-cells that exhibited their peak CS response at short latency as Type 1 (n=177), and P-cells that did not as Type 2 (n=204).

For Type 1 cells the peak CS response was at 63±1 ms latency, whereas in Type 2 cells this response was at 129±2 ms latency (Fig. 3A). We next divided the trials based on RT and found that for Type 1 cells, the CS response to target onset peaked at around 65 ms regardless of whether the movement had short or long RT. However, for Type 2 cells, the peak was later in short RT trials (127±1 ms; Wilcoxon rank sum test, peak response timing in CS-on, short RT trials, Type 1 vs. Type 2, z=-8.25 p<10^-6^), and largely absent in long RT trials (Fig. 3A). Importantly, in both groups we observed a CS rate increase that peaked before saccade onset (Fig. 3B). Thus, only the Type 1 cells exhibited a CS response to the visual event.

**Figure 3.**
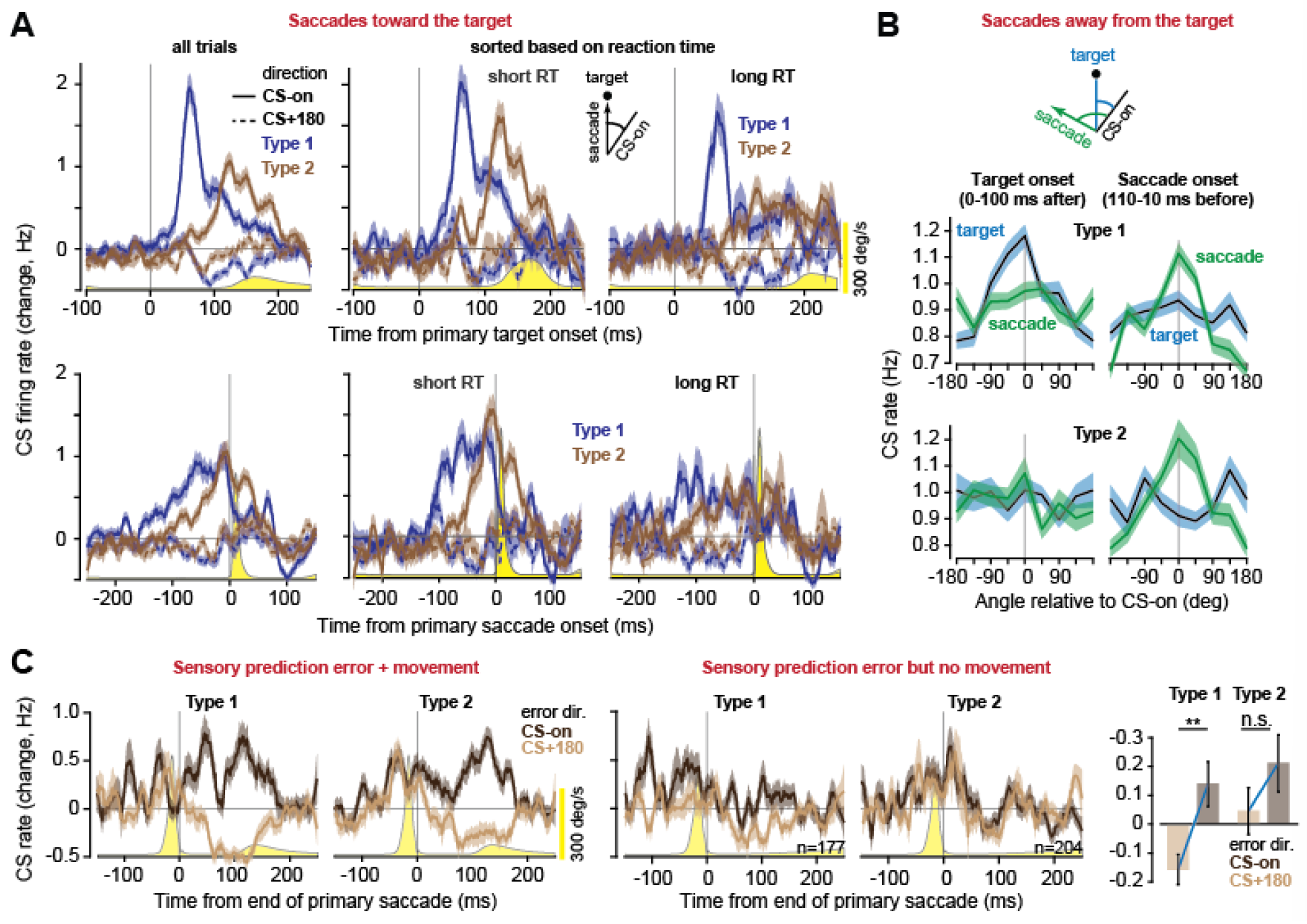
The olive reported sensory prediction errors to only Type 1 P-cells. **A**. The CS response (change from baseline) of each group of P-cells aligned to the presentation of the primary target, and to the onset of the primary saccade, for short and long reaction time (RT) trials. Only the Type 1 P-cells responded to the visual event. **B**. Trials in which the subject was presented with a visual target but chose to saccade elsewhere. The plots show the CS rate as a function of target direction, or saccade direction, with respect to CS-on. Following target onset, the CS rates in Type 1 P-cells, but not Type 2 P-cells, encoded the direction of the target. Before saccade onset, the CS rates in both groups encoded the direction of the forthcoming movement. **C**. At the end of the primary saccade, the target was not on the fovea, but moved to a random secondary location. Left plot: trials in which the subject chose to make the secondary saccade. Both P-cell types had a CS response that reported the direction of the error. Center plot: trials in which the subject chose to suppress the secondary saccade. The direction of the sensory prediction error was reported only to Type 1 P-cells. Bar plot: CS rates in the 100 ms period following end of primary saccade. Yellow trace is saccade velocity. Error bars are SEM.

We had labeled the P-cells based on their response to the primary target, but to check the robustness of this classification we considered their responses to the secondary and center targets. We found that in all cases, during the 25-100 ms period following target presentation Type 1 cells had a larger CS response than Type 2 cells (Type 1 vs. Type 2; secondary target response, z=7.70 p<10^-6^; center target response, z=11.0 p<10^-6^) (Supplementary Fig. S3).

We next focused on trials in which the subject was presented with a visual target but chose to saccade elsewhere. In Type 1 cells but not Type 2 cells, after target onset (0-100 ms period) the CS rates demonstrated a better encoding of target direction as compared to saccade direction (Fig. 3B, difference in the CS-on response, target minus saccade; Wilcoxon signed rank test; Type 1 z=3.66 p=1x10^-4^, Type 2 z=0.94 p=0.17). However, before the saccade the direction of the forthcoming movement was reflected in the CS rates of both Type 1 and Type 2 cells: at -110 to -10 ms before saccade onset in both cell types the CS rates displayed a better dissociation of saccade direction vs. target direction (difference in the CS- on response, saccade minus target; Wilcoxon signed rank test; Type 2 z=2.14 p=1x10^-2^; Type 1 z=1.96 p=0.02). A follow up analysis showed that the olivary input to the Type 2 cells better encoded saccade direction than target direction (difference in saccade CS-on and target CS-on responses, saccade onset period vs. target onset periods; Wilcoxon signed rank test z=2.47 p=7x10^-3^).

We estimated that based on our MRI of the subject’s cerebellum and the trajectory of the electrode, most of the Type 1 cells were located in the anterior part of lobule VII, whereas most of the Type 2 cells were in the posterior part of lobule VI (Supplementary Fig. S2C). Moreover, Type 2 cells had a higher baseline CS rate than Type 1 cells (Supplementary Fig. S2D, Wilcoxon rank sum test, Type 1 vs. Type 2, z=4.50 p<10^-6^).

In summary, following presentation of a visual stimulus at a random direction, at short latency the Type 1 P-cells received information from the olive regarding the direction of that stimulus. This sensory information was not sent to the Type 2 cells. However, just before saccade onset, both groups received information from the olive regarding the direction of the forthcoming movement. Thus, the olive differentiated the P-cells based on whether they would receive sensory information or not.

### The olive reported sensory prediction errors to only Type 1 cells

Following the completion of the primary saccade, the target was not on the fovea, resulting in a sensory prediction error. We used reward to assign a value to this error: in some sessions reward required a corrective movement (i.e., a secondary saccade), whereas in other sessions the subject had the option of withholding the corrective movement (in these trials reward required only the primary saccade). We labeled the trials across all sessions based on whether the animal chose to correct for the error or not.

We began with trials in which the subject chose to correct for the error, i.e., produce a secondary saccade. If the error was in direction CS-on, and the subject chose to correct it, in Type 1 cells the CS rates peaked at around 60 ms latency (Fig. 3C, left subplot). Notably, this short-latency visual response was missing in Type 2 cells, confirming what we had seen in response to the primary target (Fig. 3A). If the error was in direction CS+180, both groups exhibited CS suppression that reached a peak around 100 ms following target onset. Thus, the CS response to the secondary target was similar in timing to the response that the cells had produced to the onset of the primary target. This is consistent with the fact that both the primary and secondary targets were random and thus constituted sensory prediction errors.

Critically, if the subject chose to suppress the corrective movement (Fig. 3C, right subplot), then in the Type 2 cells the CS rates no longer encoded the error direction (0-100 ms period, Wilcoxon signed rank test; z=1.34 p=0.09). As a result, without the corrective movement the sensory prediction error was insufficient to drive the olivary inputs to Type 2 cells. In contrast, in Type 1 cells, even without the corrective movement the presence of the error was sufficient to modulate the CS rates (Fig. 3C bar plot, CS-on vs. CS-off, 0-100ms post primary saccade end, Wilcoxon signed rank test; z=2.95 p=2x10^-3^). This modulation was much weaker than when the error was followed by a movement (vs. error corrected, CS-on, 0-100ms post target onset; Wilcoxon signed rank test; z=4.64 p=2x10^-6^). That is, sensory prediction errors that did not produce a corrective movement resulted in a weak but still significant modulation of the CS rates, but in Type 1 cells only.

Thus, information regarding the sensory prediction error was sent only to Type 1 cells. Moreover, this error by itself was a weak modulator of CS rates when it was not accompanied with a movement.

### Omission of an expected sensory event suppressed CS rates in Type 1 cells but not Type 2 cells

At the conclusion of a saccade the target should be on the fovea. Here, at the end of the primary saccade, the target was not on the fovea but placed elsewhere. If the olivary input transmitted prediction errors, and not just sensorimotor events, then this omission by itself should leave its mark in the CS rates (10). Moreover, if Type 1 cells are the only ones that receive sensory information from the olive, then the omission should be reflected in their CS rates, but not in Type 2 cells.

To directly test for this idea, we needed a condition in which the saccade ended but the target was blanked. We did not have such a condition. To approximate this situation, we aligned the data based on the direction of the primary saccade, and then averaged across all secondary target locations. As a control, we considered saccades that did not experience endpoint errors, i.e., the secondary and center saccades. As a second control we estimated a 95% confidence interval on post-saccadic CS rates via bootstrapping: we replaced the direction of the primary saccade with a random direction.

In general, regardless of whether there was an error at the end of the saccade or not, following a saccade (in any direction) the CS rates were suppressed below baseline (Supplementary Fig. S4, Wilcoxon signed rank test; all direction; Type 1 z=-8.72 p<10^-6^; Type 2 z=-6.14 p<10^-6^). However, in Type 1 cells (but not Type 2) the CS suppression was tuned as a function of the direction of the primary saccade: the maximum CS suppression occurred following primary saccades that were in direction CS-on (comparison with bootstrap distribution, p<1x10^-3^).

To confirm that this was a response to the omission of the target rather than a generic post-saccade response, we checked and found that the CS response did not exhibit a directional tuning for saccades that did not experience error (Supplementary Fig. S4, no error trials, i.e., secondary and center saccades). These results suggest, but do not prove, that at the conclusion of the primary saccade, the omission of the target on the fovea was an event that by itself generated an error signal, resulting in the suppression of the CS rates. Critically, the effect of this omission was present only in the Type 1 cells.

### Simple spikes were suppressed during fixation, did not respond to target presentation, but exhibited an agonist-antagonist burst during saccades

The P-cell types differed in terms of the input that they received from the olive. Did their simple spikes differ in how they contributed to behavior? To answer this question, we turned to the SSs that we had simultaneously recorded with the CSs. We will first present the results for the entire population, then focus on the two cell types.

During fixation of the center target, the SS rates were suppressed below baseline (Fig. 4A, first row, Wilcoxon signed rank test; -50 to 50ms post target onset, CS-on: z=4.27 p=2x10^-5^; CS-off: z=4.03 p=6x10^-5^). Following presentation of the primary target, the SS rates did not exhibit a sharp change (unlike the CS response). Indeed, even in long RT trials the SS rates remained suppressed during fixation and did not appear to respond to target presentation (Fig. 4A, 2^nd^ row). Thus, target direction information was reported to the P-cells at short latency via the climbing fibers, but apparently not via the parallel fibers.

**Figure 4.**
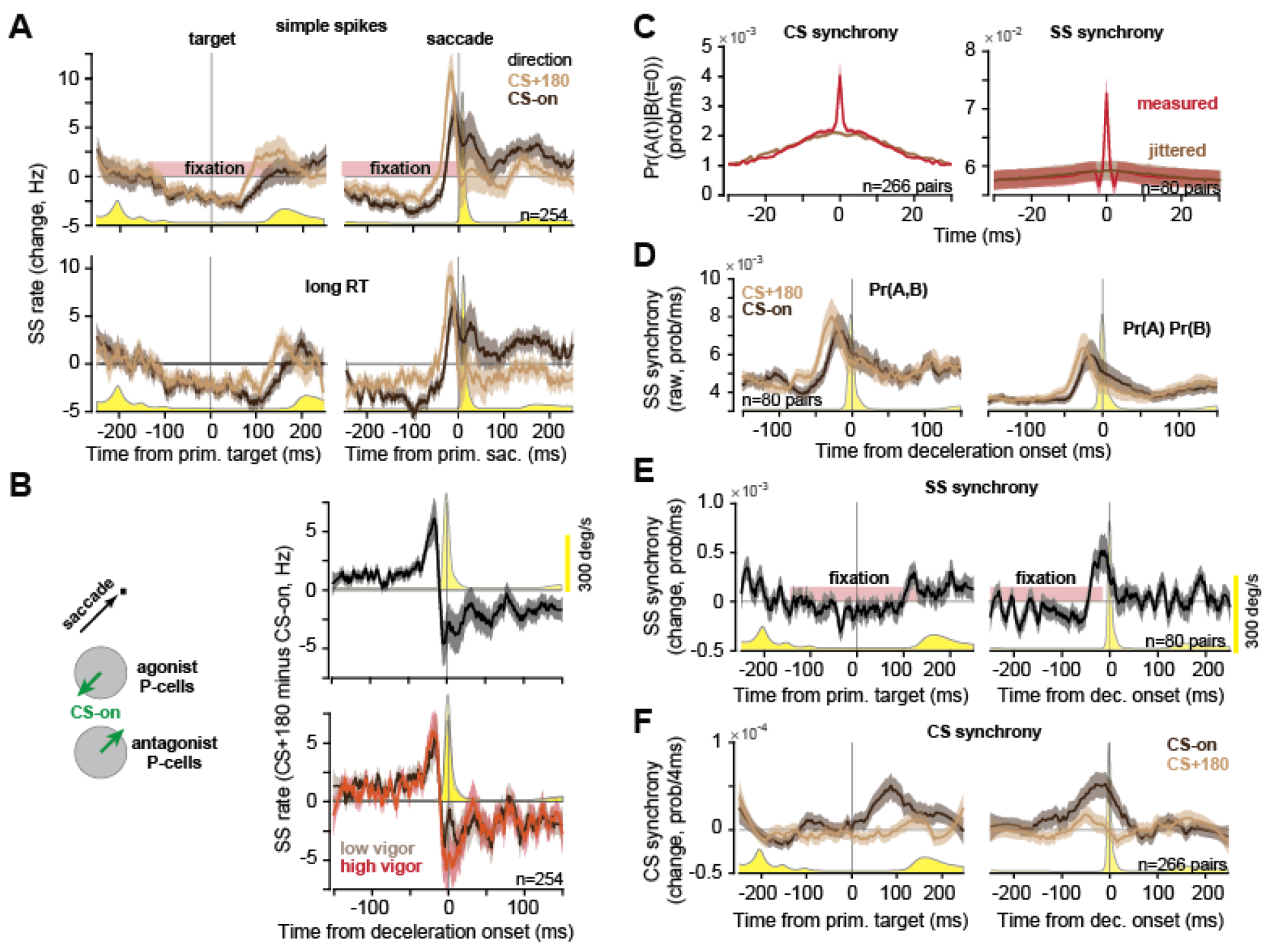
Simple spikes were suppressed during fixation and then exhibited an agonist-antagonist synchronous burst during saccades. **A**. Top row: SS rates (with respect to baseline) aligned to primary target onset and saccade onset. Bottom row: SS rates in trials that exhibited long reaction times (RT). There is little or no SS response to the visual event, but a burst before the motor event. The burst is stronger and occurs earlier for saccades in direction CS+180. **B**. SS population response. When a saccade takes place at an arbitrary direction, for one group of P-cells that saccade is in direction CS-on, while for another group the same saccade is in direction CS+180. The population response was computed as the SS rates in direction CS+180 minus CS-on. Top: all primary saccades. Bottom: primary saccades divided based on low or high peak velocity. **C**. SS and CS synchrony. The plot shows Pr(𝐴(𝑡) = 1|𝐵(0) = 1), i.e., the probability of spiking in neuron A at time t, given that neuron B spiked at time 0 (using a 1 ms time bin). Chance was estimated via spike time jittering (20 ms intervals). Error bars are 99% confidence. Time bin is 1 ms. **D**. SS raw synchrony Pr (𝐴, 𝐵) and chance synchrony Pr(𝐴) Pr(𝐵) for primary saccades. Time bin is 1 ms. **E**. SS synchrony, measured via Eq. (1). Time bin is 1 ms. **F**. CS synchrony. Time bin is 4 ms. Unless noted, the error bars are SEM. Yellow trace is eye velocity.

In contrast, near saccade onset there was a sharp increase in the SS rates. Notably, the burst was stronger and earlier if the saccade was in direction CS+180 (peak at ∼20 ms before saccade onset) as compared to direction CS-on (∼10 ms before saccade onset, Wilcoxon signed rank test, CS-off vs. CS-on, - 20ms pre target onset, z=3.56 p=2x10^-4^).

To interpret the meaning of the SS activation patterns, we used a simple model. When a saccade took place in an arbitrary direction (Fig. 4B, left panel), for one group of P-cells that saccade was in direction CS-on, while for another group that same saccade was in direction CS+180. During the movement the SSs produced by these two groups were likely to have downstream effects that opposed one another. This is because there is a relationship between the CS-on direction of a P-cell, and the direction of action signaled by the SSs of that P-cell: when the SSs of a P-cell are briefly suppressed during a saccade (because of arrival of a CS), the result is downstream motor commands that pull the eyes in direction CS-on of that P-cell (31). Similarly, when the SSs are activated via optogenetics, the result is downstream commands that pull the eyes roughly in direction CS+180 (32).

Based on these results, we imagined that when the movement was in direction CS+180 of a group of P-cells, the SSs produced by those cells were functionally the agonist of that saccade, commanding downstream forces that aided acceleration. On the other hand, when the movement was in direction CS-on, then the SSs were functionally the antagonist, commanding downstream forces that produced deceleration. In this model, the population response is measured by subtracting the SS rates in direction CS-on from the SS rates in direction CS+180. When we did this, the result unmasked a striking pattern (Fig. 4B, 1^st^ row): the net activity was a burst that preceded saccade onset in the P-cells for which the movement was in their CS+180 direction, followed by a burst that reached its peak roughly 10 ms before deceleration onset in the P-cells for which the movement was in their CS-on direction (Fig. 4B, 1^st^ row). Thus, the movement was preceded by activity in P-cells whose simple spikes likely signaled a motor command in the direction of the movement, and then just before deceleration onset, the activity switched to P-cells whose simple spikes likely signaled a motor command opposite to the direction of movement.

As the motivational state of the subject changed, so did saccade peak velocity (33). We measured peak velocity as a function of saccade amplitude in each subject and then normalized the data to compute vigor of each saccade (34, 35). For high vigor saccades, there was little change in the agonist SS burst, but there was an increase in the antagonist burst around the time that the saccade began decelerating (Fig. 4B, 2^nd^ row, 2^nd^ row, high vs. low vigor, Wilcoxon signed rank test; from -75 to -25ms of deceleration onset, z=0.55 p=0.59; from -10 to +20 ms of deceleration onset, z=-3.17 z=2x10^-3^).

In summary, the SSs were suppressed during fixation, and did not respond to target presentation. However, around 50 ms before saccade onset the SSs exhibited a burst that was earlier and stronger when the saccade was in direction CS+180, weaker and later when the saccade was in direction CS-on. The population response, measured as the SS activity in direction CS+180 minus the SS activity in direction CS-on, was an agonist burst that preceded saccade onset, followed by an antagonist burst that reached its peak just before deceleration onset.

### Both simple and complex spikes became more synchronized during saccades

Our data included n=266 pairs of P-cells in which the CSs were isolated in both cells, and n=80 pairs of P-cells in which both the CSs and SSs were isolated in all cells. To ask whether there was a significant temporal coordination among the spikes, we computed Pr(𝐴(𝑡) = 1|𝐵(0) = 1), i.e., the probability of spiking in neuron A at time t, given that neuron B spiked at time 0 (using a 1 ms time bin). Because this quantity varied with the firing rates and those rates were modulated by the task, as a control we used interval jittering (20 ms intervals) to randomly move the spikes without altering their average firing rates (36). By repeating the jittering 1000 times, we computed a 99% confidence interval (CI). This allowed us to compare the measured values with chance.

We found that if a CS occurred in one P-cell, then there was roughly 2x greater than chance probability that the second P-cell would also produce a CS during the same 1 ms time bin (Fig. 4C). This probability returned to chance levels at ±5 ms delay. When an SS occurred in one P-cell, then there was roughly 1.2x greater than chance probability that the second P-cell would also produce an SS during the same 1 ms time bin. This probability returned to chance at ±1 ms delay. Thus, the CSs and SSs were on average 2x and 1.2x more likely to be synchronous within a 1 ms time bin than expected by chance.

We next asked whether the probability of synchronous spiking was modulated by the task. To compute the probability of a synchronous event with respect to chance, we used the approach suggested in (37) and aligned the trials to target onset or movement onset and then for each time bin computed the following quantity:

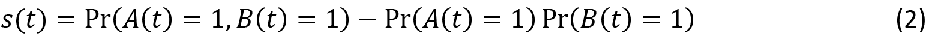

In the above equation, Pr (𝐴, 𝐵) is the “raw” probability of synchronous spiking, while Pr(𝐴) Pr(𝐵) is the expected synchrony if the two neurons were independent. Pr (𝐴) and Pr (𝐵) are the measured probability of spiking in each neuron at that time bin (reflecting the average firing rate). For this analysis, we used a 1 ms bin to label a synchronous SS event, and a 4 ms bin to label a synchronous CS event.

Thus, 𝑠(𝑡) quantified the probability of synchrony with respect to chance. To ask if there were task related changes in synchrony, for each cell pair we measured synchrony with respect to baseline, 𝑠(𝑡) −𝑠_0_, where 𝑠_0_is baseline synchrony in that cell pair (𝑠_0_is a constant measured during a task irrelevant period of the task, see Methods).

For the SSs both the raw synchrony Pr (𝐴, 𝐵) and the chance synchrony Pr(𝐴) Pr(𝐵) increased (Fig. 4D) near the onset of the primary saccade, as would be expected because of the increase in the firing rates. However, the observed synchrony rose faster than expected from chance, and thus at around 50 ms before saccade onset the quantity 𝑠(𝑡) began to rise and reached a peak just before saccade deceleration (Fig. 4E, Wilcoxon signed rank test, -20 to 0 ms period before saccade deceleration, z=2.8 p=5x10^-3^).

The CSs also exhibited task related synchrony. Following presentation of the primary target in direction CS-on, CS synchrony increased, reaching a peak at around 100 ms (Fig. 4F, Wilcoxon signed rank test, 0-200ms post target onset, z=3.39 p=7x10^-4^). Similarly, before the onset of the primary saccade, CS synchrony increased, reaching a peak just before deceleration onset (-200 to 0ms pre saccade onset, z=2.69 p=7x10^-3^).

In summary, the CSs and SSs in pairs of P-cells were on average 2x and 1.2x more likely to be synchronous than expected by chance. This temporal coordination appeared task dependent, as SS synchrony increased and reached a peak around the time that the saccade began decelerating.

### The simple spikes of Type 1 cells were suppressed during fixation

Type 1 cells received olivary information regarding target direction, but not Type 2 cells. This division of P-cells uncovered notable differences in their SS activities. We considered the fixation period, during which the subjects were required to hold the eyes at the center target. Fixation began following the conclusion of the saccade that brought the eyes to the center target (Fig. 5A, left), continued as the primary target was presented (Fig. 5A, center), and ended at the onset of the primary saccade (Fig. 5A, right). Type 1 cells exhibited greater SS suppression than Type 2 cells (Wilcoxon rank sum test; 100 ms fixation period centered on primary target onset, Type 2 > Type 1, z=3.57, p=2x10^-4^).

**Figure 5.**
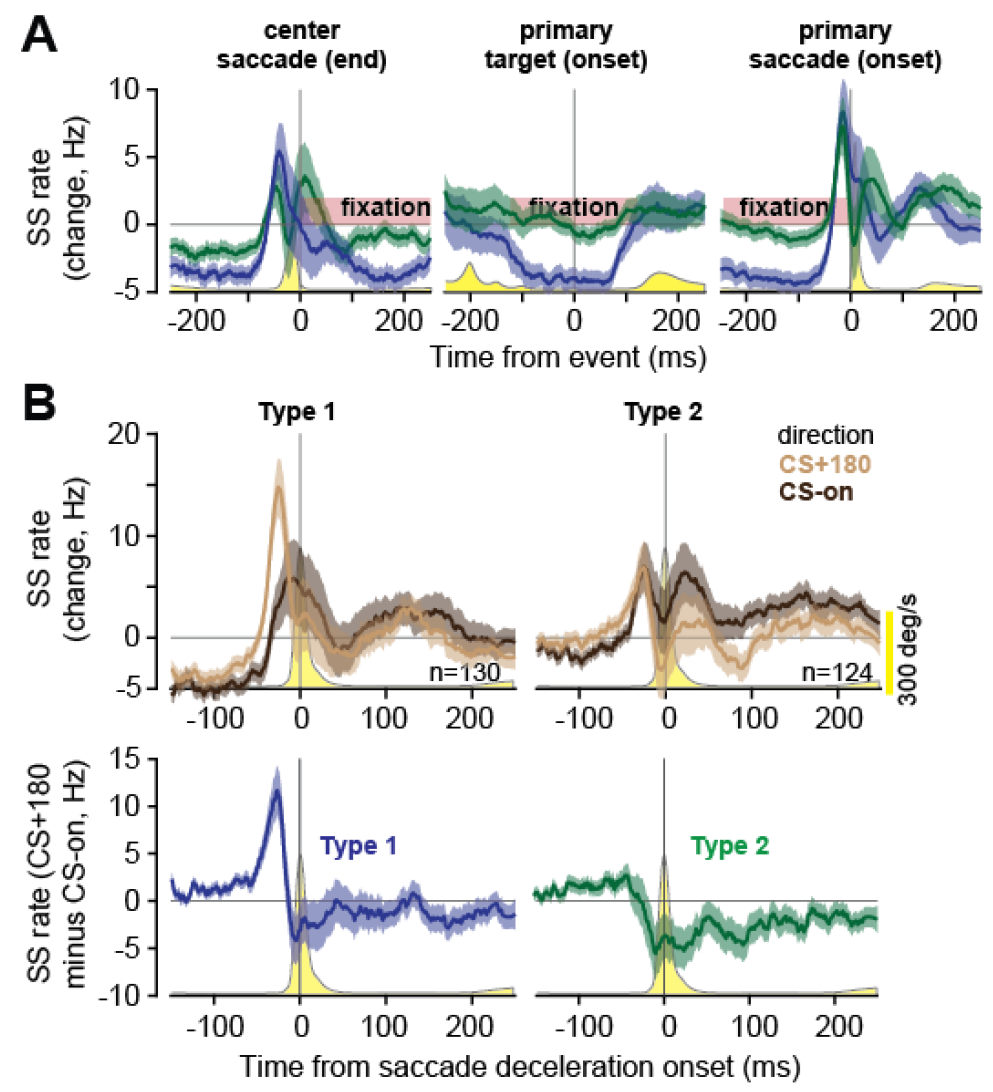
Type 1 P-cells produced an agonist SS burst while Type 2 P-cells produced an antagonist burst. **A**. SS activities in each group of P-cells during center target fixation. Data aligned to start of fixation (end of center saccade), during fixation, and end of fixation (primary saccade onset). **B**. SS rates (with respect to baseline) during the primary saccade. Top row: In Type 1 the burst was earlier and larger in direction CS+180 before saccade onset, but in Type 2 the burst was larger in direction CS-on after deceleration onset. Bottom row: population response. Positive values are interpreted as commanding forces in the direction of the saccade (agonist). Negative values are interpreted as commanding forces opposite to the direction of the saccade (antagonist). Error bars are SEM. Yellow trace is saccade velocity.

### During saccades, Type 1 cells produced SS activity that appeared to act as agonist while Type 2 cells produced antagonist SS activity

During saccades both groups exhibited an SS burst, but the burst in the Type 1 group was unimodal, whereas in the Type 2 group it was bimodal, bifurcated by a reduction in activity near peak velocity (Fig. 5B). While in the Type 1 group the early component of the burst was stronger when the saccade was in direction CS+180, in the Type 2 group the late component of the burst appeared stronger when the saccade was in direction CS-on. Critically, the population response, calculated as CS+180 minus CS-on, differed markedly in the two groups. In the Type 1 group there was a strong agonist burst that was followed by a weak antagonist burst just before deceleration onset (Fig. 5B). In contrast, in the Type 2 group the population response lacked an agonist burst entirely but exhibited a strong antagonist burst (Fig. 5B).

When saccades were reward-irrelevant, CS modulation before movement onset was dramatically reduced (∼75% smaller than reward-relevant saccades; direction CS-on, -100 to 0 ms period with respect to peak velocity, Type 1 z=7.84 p<10^-6^, Type 2 z=8.21 p<10^-6^, Supplementary Fig. S5). SS modulation was also smaller (∼50% smaller, direction CS+180, peaks during -100 to 0 ms period with respect to peak velocity, Wilcoxon rank sum test, Type 1 z=5.49 p<10^-6^, Type 2 z=4.74 p=2x10^-6^). Despite these reduced firing rates, Type 1 cells continued to produce activity in the agonist direction before saccade onset, whereas Type 2 cells produced activity in the antagonist direction around the time of saccade deceleration.

In summary, the P-cells that received olivary input regarding the visual event (Type 1) had SSs that were suppressed during fixation, then before saccade onset produced an SS burst that was consistent with an agonist for the movement. The P-cells that received olivary input regarding movement direction but not target direction (Type 2) were not suppressed during fixation and did not produce activity that acted as an agonist. Rather, these P-cells produced an SS burst that was consistent with an antagonist for the movement.

## Discussion

To better understand the information that the inferior olive transmits to the cerebellum, we designed a task in which random visual events were sometimes but not always followed by movements, and movements that were sometimes but not always directed toward the visual events. While some movements ended in error, the subject had the option of experiencing the sensory prediction error without making a corrective movement.

We found that in the oculomotor region of the vermis, at ∼60 ms latency the olive provided the P-cells with information regarding the direction of the visual target, and then just before movement onset the olive reported the direction of the planned saccade. However, the visual and motor information were not sent to all task related P-cells. Rather, the population was composed of two groups. Sensory information was sent to one group (Type 1) but not the second group (Type 2), whereas movement information was transmitted to both groups. As a result, when a saccade ended in error, but the subject chose to suppress the corrective movement, the olive reported the error direction to only the Type 1 cells. Thus, sensory prediction errors were transmitted to Type 1 cells but not Type 2 cells.

This grouping of P-cells based on their CS response unmasked fundamental differences in their SS contributions to control of behavior. During fixation, the SSs of Type 1 cells, but not Type 2 cells, were suppressed below baseline. To estimate the contributions of each group to control of saccades, we relied on the fact that a brief suppression of a P-cell’s SS produced downstream commands that pulled the eyes toward the CS-on direction of that P-cell (38). That is, the CS-on direction of a P-cell identified both a coordinate system for the olivary input, and the direction of influence of the SSs of that P-cell. As a saccade took place, for some P-cells that movement was in their direction CS-on, while for other P-cells that same movement was in their direction CS+180. Thus, the population response was CS+180 minus CS-on. When we organized the data based on this model, a pattern emerged: the P-cells first produced an SS burst that peaked before saccade onset in direction CS+180, i.e., pulling toward the target, but then just before the saccade began decelerating, they produced a more synchronized burst in direction CS-on, i.e., pulling in the opposite direction. Notably, Type 1 cells were uniquely responsible for the SS burst that promoted the movement, acting as an agonist, whereas Type 2 cells were responsible for the SS burst that ended the movement, acting as an antagonist.

### The olive conveyed both sensory and motor information, but to different groups of P-cells

The climbing fiber input to this region of the cerebellum reflects in part the input that the inferior olive receives from the superior colliculus (39–41). For example, subthreshold stimulation of a region of colliculus leads to CS production without producing a movement (42). Indeed, in our data the CS response of the P-cells had features that were similar to the response of neurons in the superior colliculus. For example, when a target is presented in the response field of a collicular neuron (in macaques), there is a bimodal response with peaks at 60 and 110 ms (43, 44). Moreover, before saccade onset, the collicular neurons exhibit a gradual buildup, with a rise that is slower and a peak that is smaller for long RT saccades (45). The bimodal response, the gradual buildup, and the RT dependent activity of neurons in the colliculus were all present in the CS response of the P-cells.

If the olivary input to this region of the cerebellum is a stochastic representation of superior colliculus activity, then the division of P-cells into two groups may be due to the physiology of the colliculus. Neurons in the intermediate layers of the colliculus respond to both visual and motor events (46), reminiscent of Type 1 P-cells. In contrast, neurons in the deeper layers respond to movements but not visual events (47, 48), reminiscent of Type 2 P-cells. Response of collicular neurons to visuomotor events depends on the value assigned to that stimulus (49): the movement related activity of collicular neurons is smaller when that movement is reward irrelevant (50). This is consistent with our observation of a weaker CS modulation for task irrelevant saccades. We conjecture that Type 1 P-cells receive input from olivary neurons that receive their projections from the intermediate layers of the colliculus, whereas Type 2 P-cells receive input from olivary neurons that receive their projections from the deeper layers of the colliculus.

### Sensory prediction errors that did not produce movements were weak modulators of the CS rates

When a sensory prediction error occurred following a movement, the olive informed the cerebellum of the direction of the error, but weakly and only to Type 1 P-cells if the subject chose not to make a corrective movement. That is, while the CS rates were strongly modulated before the onset of voluntary movements, they were weakly modulated following sensory prediction errors that did not result in a movement.

Our results provide an explanation for an interesting behavioral puzzle: motor learning is weak when the errors are merely observed but not followed by corrective movements. For example, it is possible to perform an action, observe the erroneous sensory consequences, but withhold the correction (24, 25). The resulting error produces much smaller adaptive changes in behavior as compared to when the corrective movement is permitted. Our result explain that the CS response to error is present without a corrective movement, but muted because the Type 2 cells that depend on motor information are effectively excluded from the learning process.

Does the olivary input to this part of the cerebellum encode sensory prediction errors, or just sensorimotor events? A critical clue was the response of the Type 1 P-cells. We found that when the saccade was in direction CS-on but at movement end the target was not on the fovea, the omission of the expected sensory event produced a suppression of the CS response, but only in the Type 1 cells. If the CS response was simply an encoding of sensorimotor events, e.g., a stochastic mapping from one region of the superior colliculus to the cerebellum, then we have no reason to expect that following saccade end there should be CS suppression. Presence of this suppression suggests, but does not prove, that the CS response is partly a reflection of sensory and motor events as reported from the colliculus to the olive, and partly a reflection of cerebellar inputs to the olive that might describe expected sensory consequences of movements.

### Why are the complex spikes present at movement onset?

Learning often begins with a CS response to the unexpected sensory event, but as the error is eliminated the CS response does not vanish; rather it shifts to the onset of the movement (10). As a result, in a well-practiced behavior, the CS rates as well as CS synchrony are elevated near movement onset (18).

Indeed, we observed that both the CS rates and synchrony increased when the saccades were in direction CS-on.

One possibility is that the directional information provided by the olive before movements is necessary for maintaining the directional dependence of SS rates. In this theoretical model, the directional response of P-cells to parallel fiber input during movements is continuously maintained by the olivary input that accompanies that movement. A mechanistic explanation for how the movement related CSs may sustain the SS pattern is unknown.

A second possibility is that the purpose of the CS that accompanies a movement is not to promote learning directly, but act as a stochastic perturbation to that movement. Indeed, following a CS, saccades and reaching movements are perturbed (38, 51). The idea is that for many cerebellar-dependent behaviors, the cerebellum is not afforded the luxury of a teacher (17). In this framework, life without a teacher requires a way to stochastically change the movement, then evaluate that change to determine its utility (52). Thus, one purpose of the CSs around movement onset may be to induce stochastic changes that provides the cerebellum the means to learn without a teacher.

### A role for Type 1 P-cells during holding still

Holding still is an active process, requiring precise balance and sustained activation of antagonist muscles (53). Holmes (54) noted that a major symptom of cerebellar damage was the loss of muscle tone, as manifested during the process of holding still. Indeed, a recent report demonstrated that optogenetic excitation of P-cells in vermis lobules IV/V/VI during standing resulted in postural collapse (55). Thus, the cerebellum appears to make a fundamental contribution to the process of holding still. We found that the SS rates were suppressed below baseline during reward-relevant fixations in Type 1 cells, but not when fixations were reward-irrelevant. Thus, it appears that the oculomotor region of the vermis plays an active role both during control of saccades, and during control of fixation (56, 57). Our observation may be relevant for a recently reported clinical symptom termed “fixation nystagmus” in which patients exhibit nystagmus while fixating a visual target, but not during other periods of fixation (58).

### Population coding in the cerebellum may be via a coordinate system defined by the olivary input

The cerebellum has a fundamental anatomical feature: the olive organizes the P-cells into modules such that the P-cells that have similar olivary inputs are likely to project to nearby regions in a deep cerebellar nucleus which then project to a region in the olive from which the input to the P-cells arose (59–64).

Here, we quantified similarity in the olivary inputs via directional tuning of the CS response, labeled as CS-on. While this labeling describes a feature of the input to a P-cell, it also predicts the downstream effects of that P-cell: the CS-on direction tends to be aligned with the direction of action of the downstream nucleus neuron (6, 26, 65). Indeed, during a saccade, a brief SS suppression of a P-cell pulls the eyes in the CS-on direction of that P-cell (38). Just before the onset of bilateral symmetric whisking, a CS is more likely to be present in the P-cells of medial Crus 1 (as compared to asymmetric whisking), and optogenetic stimulation of the olivary input to this region produces symmetric whisking (66). Thus, the information encoded in the olivary input to a P-cell is critical for deciphering the downstream effects of the simple spikes of that cell.

During a saccade, for one group of P-cells the movement is in direction CS-on, while for another group that same movement is in direction CS+180. The downstream effect of their SSs is likely to oppose one another. We used this logic to compute a population response and found that before saccade onset the activity in the agonist population (CS+180) dominated, but just before the saccade began decelerating, the balance shifted to the antagonist population (CS-on). Notably, as saccade vigor changed, the antagonist activity became stronger, but not the agonist activity.

De Zeeuw (26) has proposed the microzone framework in which the identity of a P-cell is determined by the CS response of that cell to sensory stimuli. In this framework, a single behavior is composed of synergistic activities in distinct microzones of P-cells, one acting as agonist, while other acts as antagonist. Although the original framework was proposed for eye blink conditioning, our results suggest that a similar framework may be relevant for control of voluntary movements.

There are several questions that remain. If the olivary input is indeed the teacher, how do the different SS patterns in the various groups emerge? Recent findings have revealed that in primates, P-cells are more likely to have two or more dendritic branches than in mice (67), raising the possibility that a P-cell in the primate cerebellum is likely to receive more than a single climbing fiber. Do these inputs encode similar regions of the sensorimotor space, or disparate regions? In a P-cell with two climbing fiber inputs, how do the downstream effects of simple spikes relate to the information encoded in their climbing fiber inputs? These questions await further investigation.

## Methods

Data were collected from two marmosets (*Callithrix Jacchus*, male and female, 350-370 g, subjects M and R, 6 years old) during a 2.5 year period. The marmosets were born and raised in a colony that Prof. Xiaoqin Wang has maintained at the Johns Hopkins School of Medicine since 1996. The procedures on the marmosets were approved by the Johns Hopkins University Animal Care and Use Committee in compliance with the guidelines of the United States National Institutes of Health.

### Data acquisition

Following recovery from head-post implantation surgery, the animals were trained to make saccades to visual targets and rewarded with a mixture of applesauce and lab diet (27). Visual targets were presented on an LCD screen (Curved MSI 32” 144 Hz - model AG32CQ) while binocular eye movements were tracked using an EyeLink-1000 eye tracking system (SR Research, USA). Timing of target presentations on the video screen was measured using a photo diode.

We performed MRI and CT imaging on each animal and used the imaging data to design an alignment system that defined trajectories from the burr hole to various locations in the cerebellar vermis (27), including points in lobule VI and VII. We used a piezoelectric, high precision microdrive (0.5 micron resolution) with an integrated absolute encoder (M3-LA-3.4-15 Linear smart stage, New Scale Technologies) to advance the electrode.

We recorded from the cerebellum using quartz insulated 4 fiber (tetrode) or 7 fiber (heptode) metal core (platinum/tungsten 95/05) electrodes (Thomas Recording), and 64 channel checkerboard or linear high density silicon probes (M1 and M2 probes, Cambridge Neurotech). We connected each electrode to a 32 or 64 channel head stage amplifier and digitizer (RHD2132 and RHD2164, Intan Technologies, USA), and then connected the head stage to a communication system (RHD2000 Evaluation Board, Intan Technologies, USA). Data were sampled at 30 kHz and band-pass filtered (2.5 - 7.6 kHz). We used OpenEphys (68) for electrophysiology data acquisition, and then used P-sort (69) to identify the simple and complex spikes in the heptodes and tetrodes recordings, and Kilosort and Phi (70) to identify the spikes for the silicon probes.

Our data was composed of 381 P-cells that were identified because of the presence of CSs. Among these neurons, we also had excellent isolation of SSs in 254 P-cells. Included in this data were 257 P-cells in the session in which the corrective saccade was required, and from 124 P-cells in the session in which the corrective saccade was optional. The data for the session in which the corrective saccade was required were partially analyzed in a previous report (30).

### Behavioral protocol

Each trial began with fixation of a center target for 200 ms (Fig. 1A), after which a primary target (0.5x0.5 deg square) appeared at one of 8 randomly selected directions at a distance of 5-6.5 deg (50 cells were recorded during sessions with 4 random directions). The onset of the primary target coincided with presentation of a distinct tone. As the subject made a saccade to this primary target, that target was erased, and a secondary target was presented at a distance of 2-2.5 deg, also at one of 8 randomly selected directions (4 directions for 50 cells). To assign value to this erroneous outcome, in 51% of the sessions reward required both the primary and the corrective saccades, whereas in the remaining sessions reward required only the primary saccade. If reward did not require the corrective saccade, the secondary target was presented only briefly, returning to the primary location following ∼120 ms (adjusted during the recording session so that around 40% of the trials would produce a corrective saccade). Following the presentation of reward, and the center target was redisplayed.

### Data analysis

All saccades, regardless of whether they were instructed by presentation of a visual target or not, were identified using a velocity threshold. Saccades to primary, secondary, and central targets were labeled as targeted saccades, while all remaining saccades were labeled as task irrelevant. For the targeted saccades we computed the neurophysiological response to the presentation of the visual stimulus, as well as the response to the onset of the saccade. To compute the visual response to the secondary target, we aligned the data to the offset of the preceding primary saccade. For task irrelevant saccades, there were no visual targets and thus the response was aligned only to saccade onset.

The firing rates, as well as synchrony, were measured with respect to baseline. For SS rates and SS synchrony, baseline was defined as a period in which the subject was not performing a saccade and was not fixating the center target. We chose this period to be 200 to 100 ms before the onset of task irrelevant saccades during which time the subject was fixating a random location. For complex spikes, baseline rates were computed by dividing the total number of spikes by the duration of the entire recording. Instantaneous firing rates were calculated from peri-event time histograms with 1 ms bin size. We used a Savitzky–Golay filter (2nd order, 31 datapoints) to smooth the traces for visualization purposes.

CS directional tuning was computed by measuring the CS firing rates following target onset as a function of target angle with respect to the actual position of the eyes. We counted the number of CSs after target onset up to saccade onset or a fixed 200 ms window, whichever happened first. Dividing the spike count by the duration of time resulted in the CS firing rate as a function of target direction. A vector sum across the directions produced a vector that we labeled CS-on.

When the target was in direction CS-on, each P-cell produced a CS response that exhibited a peak at a specific time with respect to target onset (Fig. 2A). Across the population, the distribution of CS timing appeared bimodal (Supplementary Fig. S2A). We performed statistical testing to quantify whether the measured distribution was composed of one, two, or three distinct groups of neurons (Supplementary Fig. S2B). To perform this test, for each cell we measured the timing of the peak CS response, labeled it with a random variable 𝑌, and then represented the resulting distribution as a mixture of gamma functions (Eq. 1), where the number of neuronal groups, and their parameters, were unknown. We fit Eq. (1) to a mixture of 1, 2, or 3 groups of neurons and for each produced a goodness of fit, and then corrected for model complexity using AIC and BIC (Supplementary Fig. S2B). The results revealed that the distribution was best fit with a model in which there were two distinct groups of neurons, labeled as Type 1 and Type 2.

To compute properties of the CS response (i.e., timing of the maximum and minimum), we used bootstrapping, randomly sampling from our population to produce a pseudo population that consisted of 70% of the neurons. We repeated this procedure 50 times to produce error bars on our estimated timing. This procedure produced a distribution associated when the peak response occurred when the target was in direction CS-on, and when the minimum response occurred when the target was in direction CS+180.

To compute SS population response during saccades, we began by computing the change in SS firing rate of each P-cell with respect to its baseline. Next, we labeled each saccade by measuring its direction of movement with respect to the CS-on of the P-cell. Finally, we summed the activities in all P-cells (i.e., changes with respect to baseline) for saccades in direction CS-on, CS+45, etc., using a bin size of ±25 deg.

### Synchrony analysis

With multi-channel electrodes, the activity of one neuron can easily be present on multiple channels, thus giving the illusion that the two channels are picking up two distinct neurons. To guard against this, we waveform triggered the data recorded by channel A by the spikes recorded on channel B. This identified the waveform of the neuron recorded by channel B on the spike recorded on channel A. We compared this cross-channel triggered waveform with the within channel triggered waveform generated by the spikes recorded by channel A. The cross-channel triggered waveform must produce a different cluster of spikes in A than the main neuron isolated by A. If there were spikes that occurred within 1 ms of each other on channels A and B, we used these coincident-spike events to trigger the waveform in A. The spikes in A that were identified to be coincident with B should look approximately the same as the non-coincident spikes in A. Examples of this approach are provided in Sedaghat-Nejad et al. (27).

Second, we jittered the spike timing for each pair of cells to produce a measure of chance level synchrony. We followed the interval jittering procedures described in (36): the entire recording was divided into 20 ms intervals and spikes were moved within their designated interval. We jittered each recording 50 times and computed the chance level probabilities.

To compute synchrony, we aligned the trials to visual or motor events and then at each time bin computed the quantity defined in Eq. (1) (37). This measure quantified the probability of coincident spikes with respect to that expected from two independent neurons. We used a 1 ms time bin to compute SS synchrony, and a 4 ms time bin to compute CS synchrony. Next, to measure task related change in synchrony, for each cell pair we measured synchrony with respect to its baseline: 𝑠(𝑡) − 𝑠_0_, where 𝑠_0_ is the baseline synchrony for that cell pair as measured during a task irrelevant period of the task (same period for which baseline SS rates were measured, 200-100 ms period before task irrelevant saccades, i.e., fixation of a random location).

### Statistics

To statistically analyze the population data, we computed p-values using a non-parametric method: Wilcoxon rank sum.

---

**Supplementary Fig. S1.**
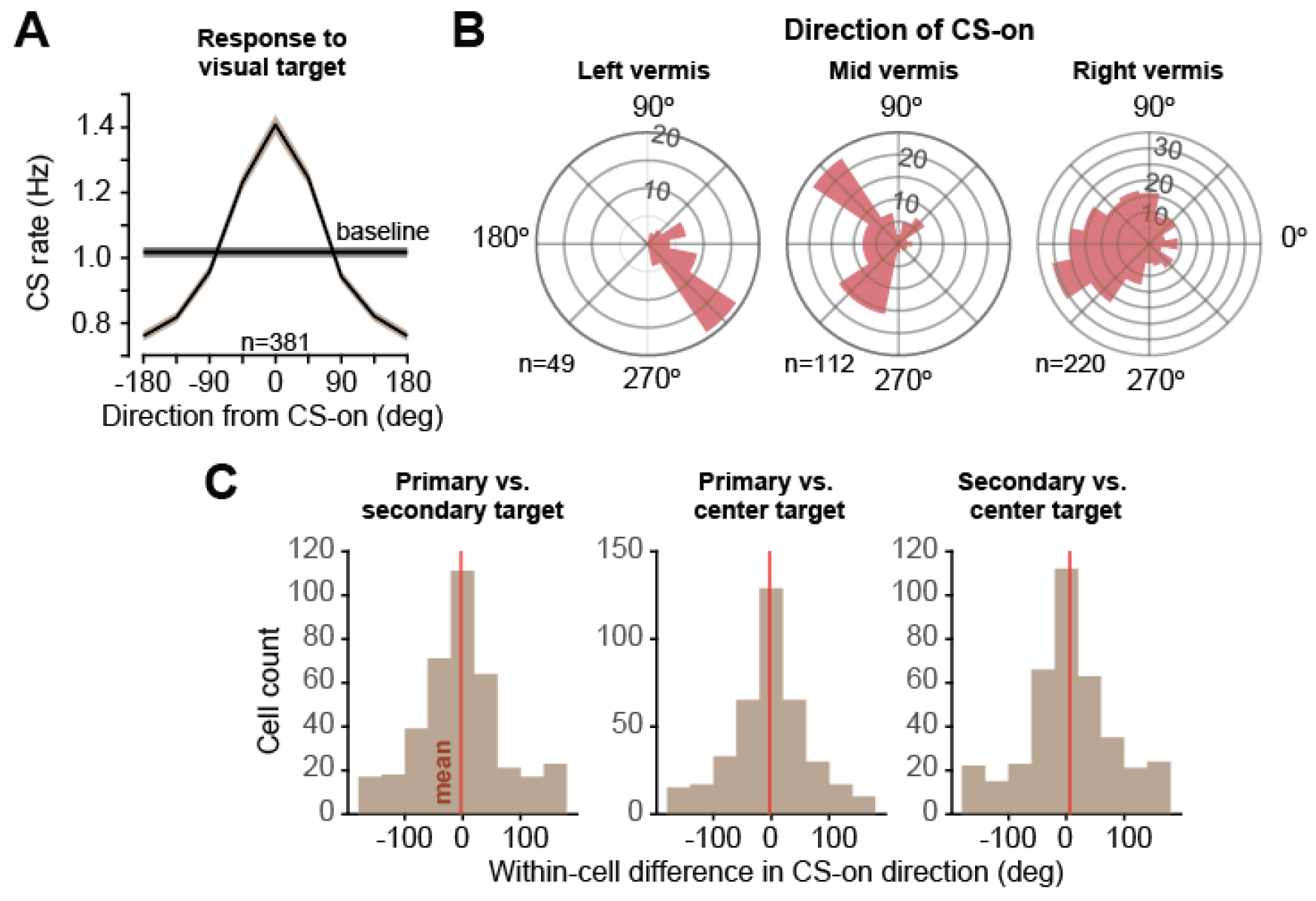
The CS response remained consistent across various targets but changed with location of the P-cell in the vermis. **A**. Average CS firing rate following the presentation of the primary target (0-200 ms period, or until saccade onset, whichever period was shorter), as a function of target direction, aligned to direction CS-on. Error bars are SEM. **B**. Distribution of CS-on as a function of location of recording in the vermis. Bin size is 15°, and the wedge at each bin represents the number of P-cells with that CS-on. P-cells tended to have a CS-on toward the contralateral hemifield. **C**. Within cell difference between CS-on directions as computed following the onset of the primary target, the secondary target, and the center target. We found no systematic differences in the estimate of CS-on between various types of targets, and thus combined the response for all targets to compute the CS-on of each P-cell.

**Supplementary Fig. S2.**
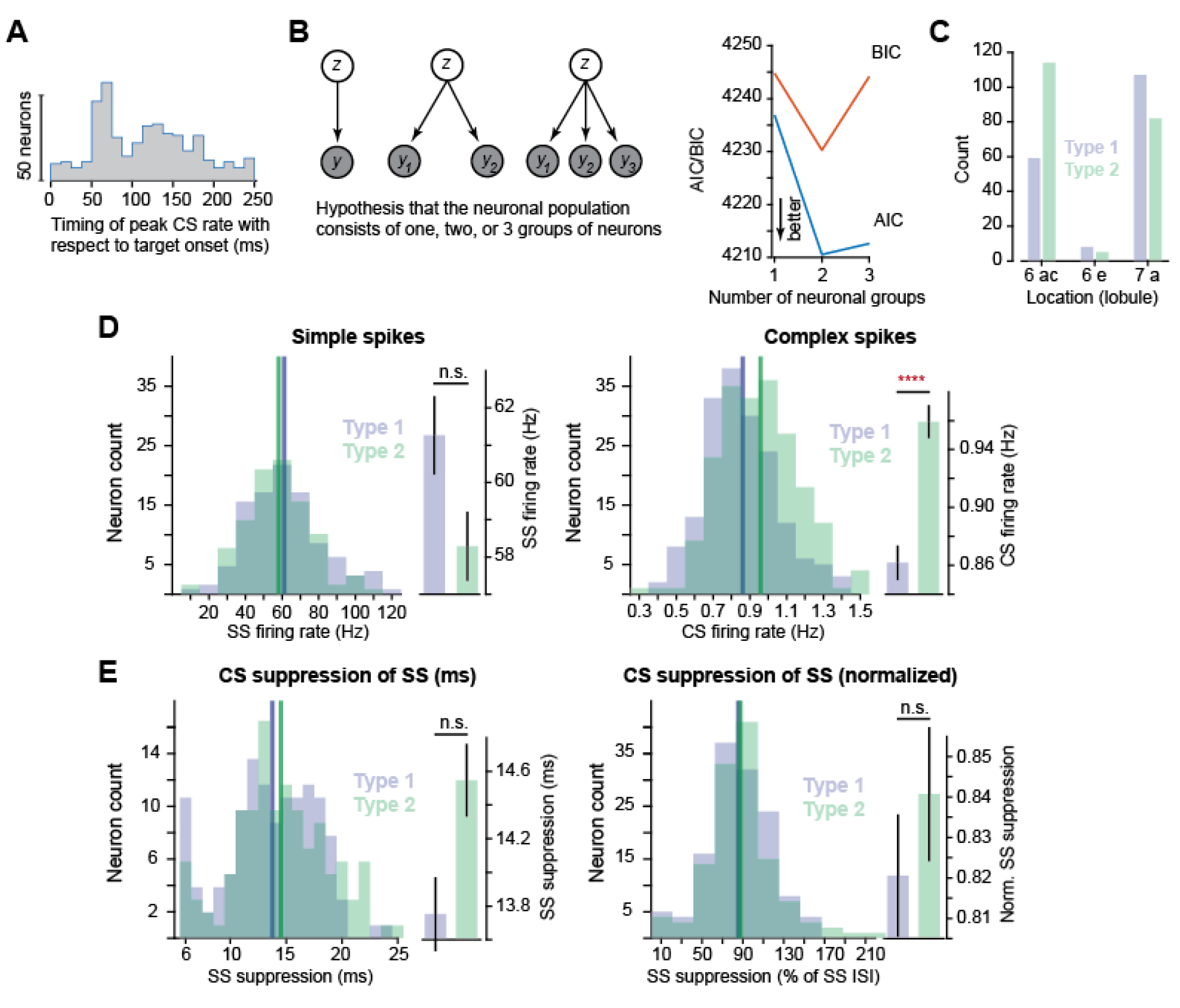
P-cells organized into two groups based on the information in their olivary input. **A**. Across neurons, the distribution of timing of the peak CS response following onset of the primary target appeared bimodal, suggesting the possibility of two distinct group of neurons. **B**. Hypothesis testing, asking whether the CS response (data in part A) indicated presence of one, two, or more groups of neurons. Models were fitted to Eq. (1). **C.** AIC and BIC measures of model fit (Eq. 1) to the distribution in part A. The data is composed of two groups of neurons. C. Approximate anatomical location of each type of P-cell in the vermis. Most Type 1 cells are in lobule 7, whereas most Type 2 cells are in lobule 6. **D**. The baseline simple and complex spike rates for each P-cell. The CS rates were significantly higher for Type 2 cells. E. Duration of SS suppression for each P-cell. The normalized measure is SS suppression duration as a function of baseline SS inter-spike interval. Error bars are SEM.

**Supplementary Fig. S3.**
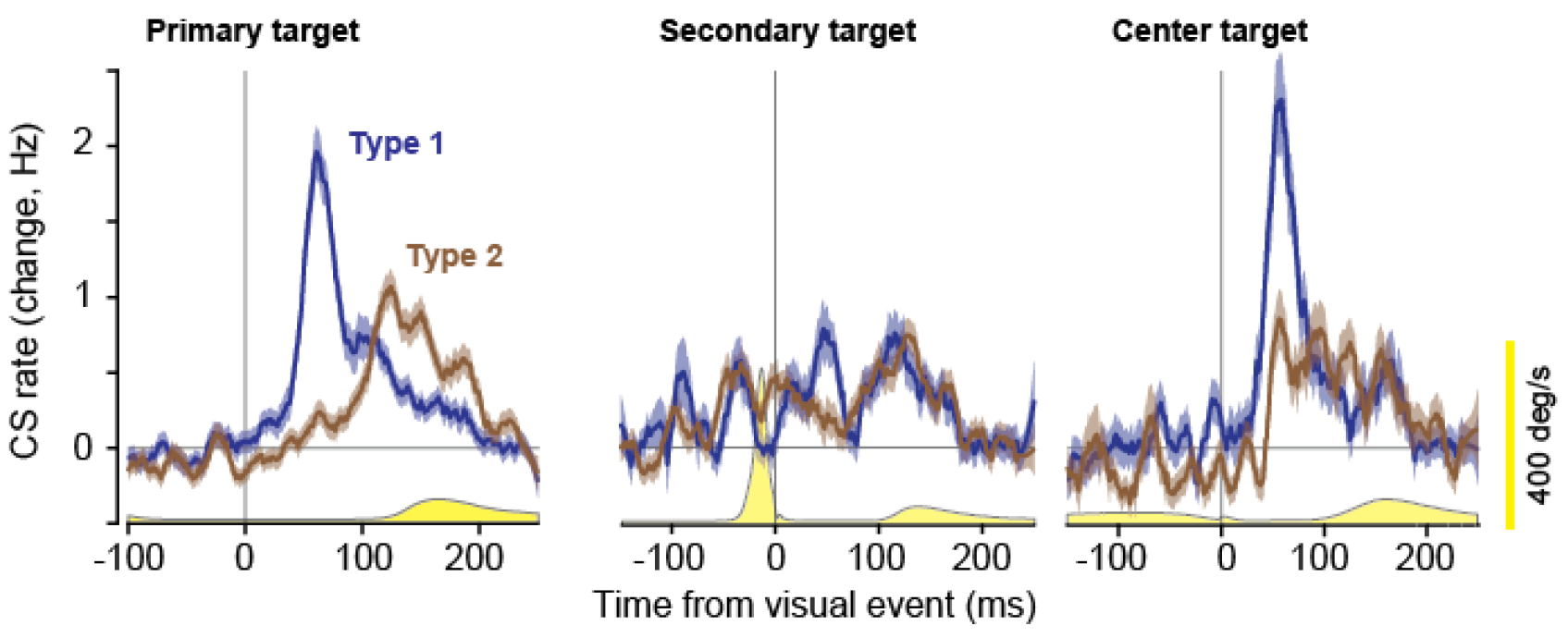
The CS response of Type 1 and Type 2 P-cells in direction CS-on following the presentation of various visual targets. The primary target was presented at a random location while the subject gazed at the center fixation. The secondary target was presented at a random location following completion of the primary saccade. The center target was presented at the center location following completion of the trial. In all cases, Type 1 cells exhibited a higher CS response (with respect to Type 2) at short latency (less than 100 ms) following target onset. Yellow traces are eye velocity.

**Supplementary Figure S4.**
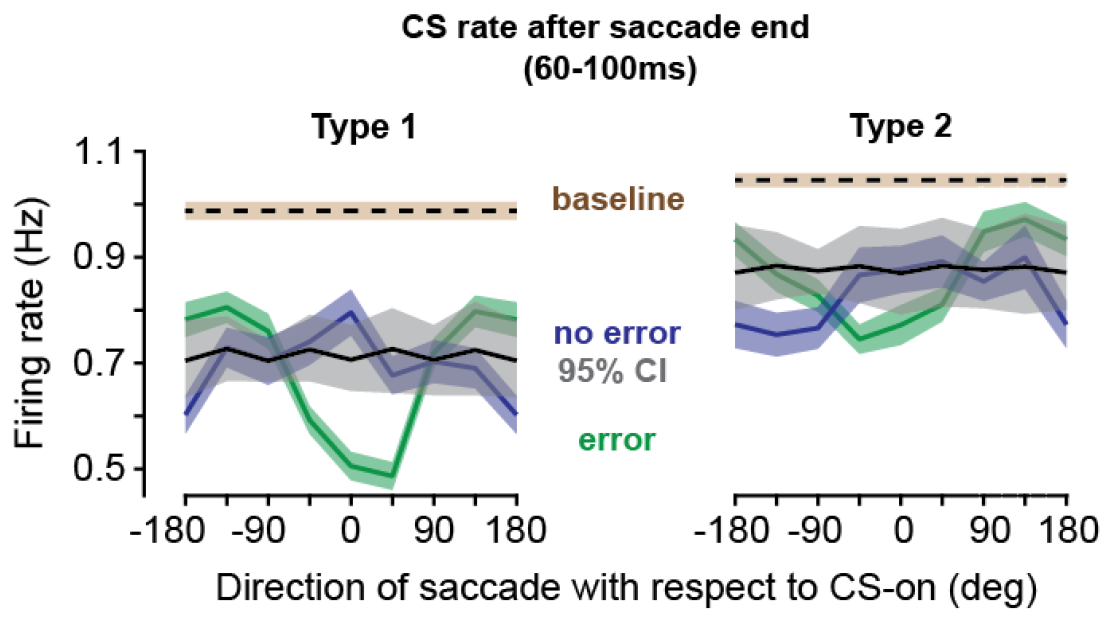
Omission of an expected sensory event suppressed CS rates in Type 1 cells but not Type 2 cells. The CS rates in the 60-100ms following completion of the saccade as a function of the direction of that saccade. Data are shown for Type 1 and Type 2 cells, for saccades that experienced endpoint error (primary saccades, green), and saccades that did not experience endpoint error (secondary and center saccades, blue). Baseline CS rates are indicated in brown. 95% confidence interval are also indicated. The 95% CI was generated by removing direction labels for the primary saccades. Error bars are SEM.

**Supplementary Figure S5.**
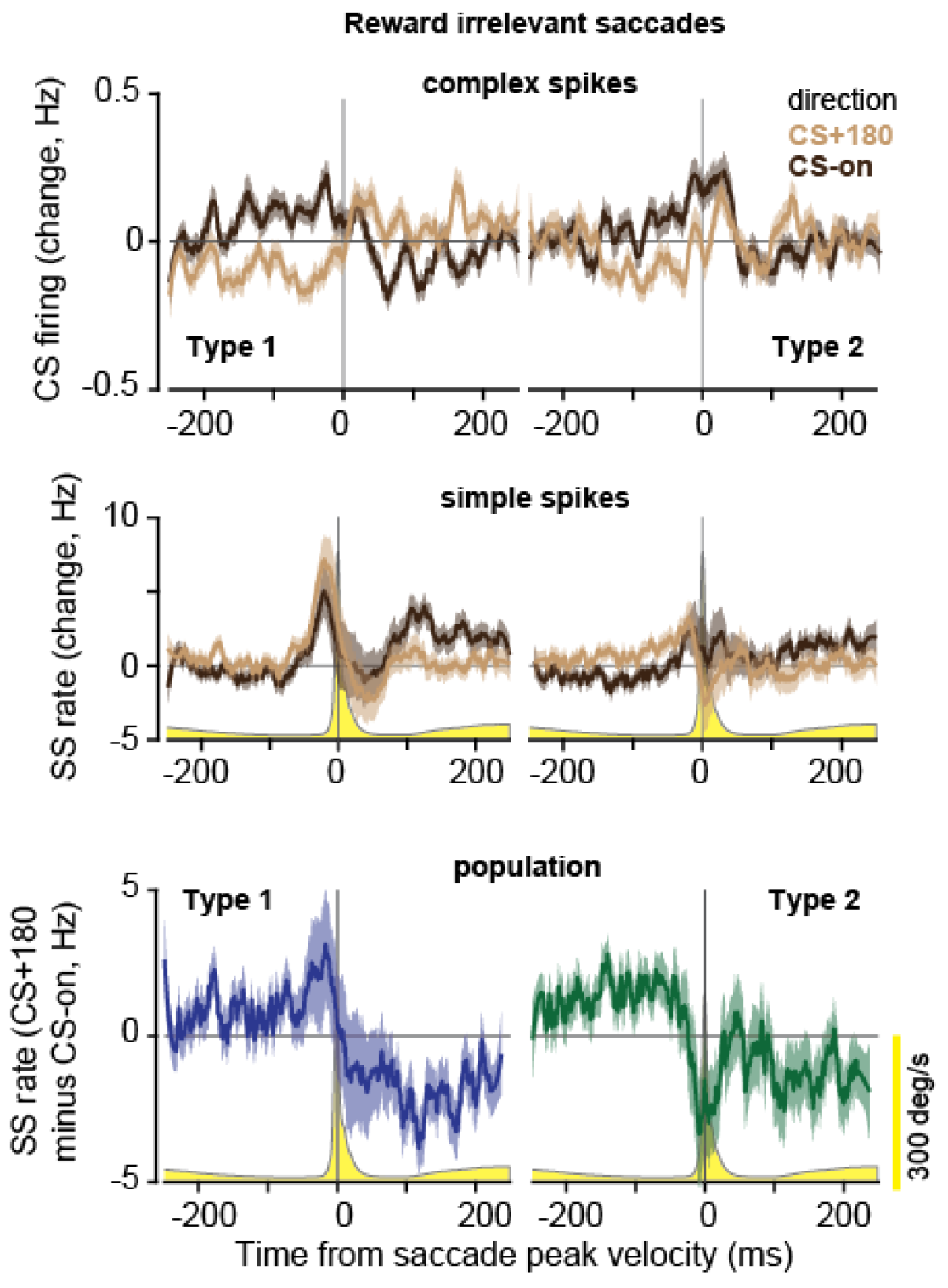
Complex and simple spike properties of reward-irrelevant saccades. The CS and SS properties of Type 1 and Type 2 saccades are illustrated. The CS response before saccade onset is roughly 75% smaller than reward relevant saccades. The SS response is also smaller, but the general agonist-antagonist pattern appears conserved in Type 1 and Type 2 cells.

